# A Novel Mouse Model Demonstrates In Vivo Replenishment of Central Nervous System Pericytes After Successful Acute Ablation

**DOI:** 10.1101/2024.09.27.614665

**Authors:** Dila Atak, Erdost Yıldız, Esra Özkan, Mohammadreza Yousefi, Ayşe Özkan, Aysu Bilge Yılmaz, Ali Burak Kızılırmak, Iman Asaad Alnajjar, Çiçek Kanar, Zeynep Lal Caan, Şakir Ümit Zeybek, Cem İsmail Küçükali, Erdem Tüzün, Yasemin Gürsoy-Özdemir, Atay Vural

## Abstract

Central nervous system (CNS) pericytes play crucial roles in vascular development and blood-brain barrier maturation during prenatal development, as well as in regulating cerebral blood flow in adults. They have also been implicated in the pathogenesis of numerous neurological disorders. However, the behavior of pericytes in the adult brain after injury remains poorly understood, partly due to limitations in existing pericyte ablation models. To investigate pericyte responses following acute ablation, we developed a tamoxifen-inducible pericyte ablation model by crossing *PDGFRβ-P2A-CreER*^*T2*^ and *Rosa26-DTA176* transgenic mouse lines. Using this model, we studied the effects of different tamoxifen doses and conducted histological examinations 15 and 60 days post-injection to assess both short- and long-term impacts of pericyte ablation. Our results demonstrate that a low dose of tamoxifen effectively ablates CNS pericytes in mice without reducing survival or causing significant systemic side effects, such as weight loss. Additionally, we found that the extent of pericyte depletion varies between the cortex and the spinal cord’s gray and white matter regions. Importantly, both pericyte coverage and numbers increased in the weeks following acute ablation, indicating the regenerative capacity of CNS pericytes in vivo. This model offers a valuable tool for future studies on the role of pericytes in neurological disorders, overcoming the limitations of constitutive pericyte ablation models.

## INTRODUCTION

Pericytes are mural cells located on the abluminal side of microvessels^1^. The pericyte/endothelial cell ratio is highest in the retina and brain, emphasizing their critical role in these regions^2-4^. In the healthy central nervous system (CNS), pericytes are pivotal for maturation of the blood-brain barrier (BBB)^5,6^, regulation of blood flow^7,8^, and angiogenesis^9,10^. Previous research, including our own, has demonstrated that pericytes are implicated in the pathogenesis of various neurological disorders, such as stroke^11,12^, multiple sclerosis^13-15^, Alzheimer’s disease^16^ and CNS trauma^17,18^. Under pathological conditions, pericyte deficiency or dysfunction can lead to microcirculation deficits^11,19,20^, BBB leakage^21,22^, and can contribute to scar formation^18^ and neuroinflammation^23^.

Pericyte ablation models are frequently used to study the contribution of pericytes to disease pathogenesis. Previous constitutional pericyte ablation models rely on the genetic manipulation of critical molecules for pericyte development. One of the earliest models involves a complete disruption of the *Pdgfβ* gene, which resulted in embryonic lethality due to severe vascular abnormalities^24^. Later, heterozygous mice (*Pdgfβ*^+/-^) were shown to exhibit partial pericyte loss and capillary irregularities such as microaneurysm, cylindrical dilations, and enhanced endothelial cellularity^25^. When *Pdgfβ* was deleted specifically in endothelial cells, mice survived into adulthood but exhibited inconsistent pericyte coverage reduction and persistent vascular abnormalities^26^. Another approach to generate mice with partial pericyte depletion was to disrupt the retention motif of PDGFβ^27^, which resulted in BBB breakdown and increased leukocyte adhesion molecule expression, highlighting the role of pericytes in vascular integrity^6,14^. The alternative strategy to deplete pericytes is to target the *Pdgfrβ* gene, and with this strategy, complete knockouts (*Pdgfrβ*^-/-^) also experienced embryonic lethality, while heterozygous mice (*Pdgfrβ*^+/-^) showed partial pericyte deficiency and associated vascular and neurodegenerative changes^22,28,29^. In another model, point mutations were introduced into the PDGFRβ receptor that resulted in partial pericyte loss and led to increased BBB permeability and disrupted blood flow^28,30^. The limitation of these constitutional pericyte ablation modes is that they lead to developmental defects to a certain degree, hindering the accurate assessment of pericyte roles in CNS pathophysiology in adults. Therefore, there is a need for reproducible, well-characterized, and easily accessible models that allow for inducible acute pericyte ablation.

To overcome the limitations of conventional models, Nikolakopoulou et al. developed a pericyte-specific, inducible ablation model using a double-promoter approach targeting *Pdgfrβ* and *Cspg4*^31^. In this model, mice were crossed with iDTR mice, and tamoxifen was administered to the offspring, leading to the expression of the diphtheria toxin receptor in cells carrying the *Pdgfrβ* and *Cspg4* promoters. Subsequent injection of diphtheria toxin resulted in a reduction of pericytes. This model effectively demonstrated blood-brain barrier breakdown and neurovascular uncoupling in adult mice following pericyte ablation, which was associated with neurodegeneration. Although this model has been instrumental in studying CNS pericytes, it has several limitations. It requires using specific *Pdgfrβ* and *Cspg4* double-promoter mice, which are not commercially available and primarily specific to CNS pericytes. Additionally, the model necessitates a 10-day tamoxifen administration followed by a 10-day diphtheria toxin administration, which may cause systemic adverse effects^32,33^ and also may not be compatible with experiments requiring the administration of other compounds, such as in experimental autoimmune encephalomyelitis. Another approach, used by Vazquez-Liebanas et al., involved developing an endothelial-specific, tamoxifen-inducible deletion of *Pdgfβ* by crossing *Pdgfβ*^*flox*/*flox*^ or *Pdgfβ*^*flox*/-^mice with *Cdh5*(PAC)-CreERT2 mice. In the 2-month-old adult progeny, they imposed tamoxifen-inducible Pdgfβ deletion, which eventually caused a slowly progressive pericyte loss when they reached the age of 12–18 months. Using this model, the authors confirmed the role of pericytes in maintaining BBB and preventing its selective permeability. They additionally showed that adult-induced loss of PDGFβ does not lead to vessel dilation, impaired arterio-venous zonation, or the formation of microvascular calcifications, unlike observed in the constitutive developmental PDGFβ loss of function animal models^34^. However, this model is unsuitable for studying acute pericyte ablation’s effects, particularly in younger mice.

Here, we aimed to develop and characterize a novel tamoxifen-inducible acute pericyte

## MATERIALS AND METHODS

### Animals

All procedures of this study were approved by the Koç University Ethics Committee (no: 2019.HADYEK.021). The mice were kept in a regulated facility at 22 ± 2 °C with a 12-hour light/dark cycle to ensure optimal acclimatization before the experiments began. The animals were housed in groups of no more than five per cage and had continuous access to food and water.

*PDGFRβ-P2A-CreER*^*T2*^ (JAX Mice, 030201), *Rosa26-DTA176* (JAX Mice, 010527) transgenic mice, and wild-type (WT) C57BL/6J mice were used for this study. *PDGFRβ-P2A-CreER*^*T2*^ line was developed by Cuervo et.al^35^. The P2A gene allows ‘‘ribosome skipping” between the *Pdgfrβ* and *CreER*^*T2*^ coding sequences resulting in the production of related proteins at similar expression levels. *Rosa26-DTA176* line was developed by Wu et. al.^36^. We chose this line because DTA176 is an attenuated form of fragment A of the diphtheria toxin (DTA). Once inside a cell, one molecule of DTA is sufficient to kill the target cell, whereas DTA176 is toxic at approximately 100-200 molecules per cell. This approach mitigates potential issues related to the leaky expression of diphtheria toxin before Cre-mediated activation^36^. To segregate mice into ablation model to overcome these limitations. To achieve this, we crossed the *PDGFRβ-P2A-CreER*^*T2*^ and *Rosa26-DTA176* transgenic mouse lines and tested various tamoxifen doses to determine the optimal dose that induces effective pericyte ablation without causing significant systemic adverse effects. Our study establishes that acute pericyte ablation can be induced in adult mice without causing significant systemic adverse effects. Using this model, we showed that CNS pericytes are replenished *in vivo* after the acute pericyte ablation period is over. The model described here can be used in future studies to assess the role of pericytes in various disorders in adult mice in a customarily developed vascular system.

appropriate experimental cohorts, after mating of the *PDGFRβ-P2A-CreER*^*T2*^ and *Rosa26-DTA176* lines, the offspring was genotyped to identify the inducible CreER^+^ and non-inducible CreER^-^ transgenic mice. The primers used, and the details of the touch-down PCR are given in Supplementary Table 1 and Supplementary Table 2.

### Tamoxifen Administration for Induction of Pericyte Ablation

Tamoxifen was prepared as a 10 mg/ml solution in corn oil and administered intraperitoneally to mice at 100 mg/kg/day over two (2X group), three (3X group), or five (5X group) days depending on the experimental design. Tamoxifen treatment initiated the CreER-mediated genetic recombination process only in the CreER^+^ mice, selectively ablating *Pdgfrβ*-expressing pericytes through the induced expression of the DTA176.

### Experimental Design

The experimental setup included both male and female mice aged between 8-16 weeks, and the studied phenotypes did not differ between sexes. Mice were systematically divided into CreER^+^ and CreER^-^ groups for detailed comparison. Within these divisions, mice were further categorized into subgroups based on the duration of tamoxifen treatment (2, 3, or 5 days). Follow-ups on these mice were conducted over 15 (acute phase) or 60 days (chronic phase), with body weight tracking and clinical assessments. After the completion of the experiments, animals were euthanized. First, animals were anesthetized with a cocktail of 100 mg/ml ketamine and 20 mg/ml xylazine (0.1 ml/10 mg body weight) injection. Transcardial perfusion was carried out using ice-cold phosphate-buffered saline (PBS), followed by fixation with 4% paraformaldehyde (PFA) in PBS, and tissues were extracted for histological and molecular analyses of the brain and spinal cord.

### Immunohistochemistry

After fixation of tissues in 4% PFA at 4°C for 12 hours, brain and cervical, thoracic, lumbar, and sacral segments of the spinal cord tissues were cryoprotected using 10%, 20%, and 30% sucrose solutions until they reached equilibrium. For cryosectioning, tissues were embedded in the optimal cutting temperature compound (OCT) and rapidly frozen using liquid nitrogen vapor. Cryosections of the brain were prepared at 30 microns thickness in a free-floating manner, whereas spinal cord tissues were sectioned at 20 microns on slides. Four serial sections, spaced 100 microns apart, were collected per sample. For immunofluorescence, sections underwent blocking and permeabilization with a solution containing 5% normal goat serum (NGS), 2% bovine serum albumin (BSA), and 0.1% Triton X-100 in Dulbecco’s PBS (DPBS) for one hour at room temperature. This was followed by overnight incubation at 4°C with primary antibody against CD13 (Bio-Rad, MCA2183) at a 1:100 dilution in the blocking solution. After primary antibody incubation, sections were rinsed in DPBS and then incubated with Alexa Fluor 647-labeled anti-rabbit IgG (Abcam, ab150159) and Dylight 488-conjugated tomato lectin (Vector Laboratories, DL-1174), at 1:200 dilution for 2 hours at room temperature to visualize pericytes and vasculature, respectively. The stained sections were then mounted using a PBS-glycerol mixture containing Hoechst stain, and coverslipped for imaging. All staining procedures were performed in batches that included both the experimental and control groups to minimize technical variability during analysis. Imaging was performed on a Leica DMI8 SP8 Confocal Microscope, capturing 6-8 random cortical areas from non-adjacent brain sections approximately 100 microns apart at 20X magnification and 8-12 areas across white and gray matter at 40X magnification from different spinal cord segments.

### Image Analysis

For pericyte coverage analysis, 10-micron thick z-stack images were obtained with a 2-micron step size with confocal microscope. Then, maximum intensity projection (MIP) images were obtained and transformed into 8-bit pictures for each channel. Following Gaussian Filter (σ = 2), pictures were masked with triangle thresholding. Particles smaller than 50 pixel^2^ in size were eliminated to exclude nonspecific staining. Finally, the pericyte coverage was calculated by dividing the integrated density of the CD13 signal by the integrated density of the tomato-lectin signal. A custom MATLAB code was written to automate these processes and to avoid biased manual adjustments.

Pericytes were manually counted using ImageJ on the same MIP images. CD13-positive signals that overlapped with a Hoechst-stained nucleus (blue) and were located on a tomato lectin-stained vessel (green) were identified as pericytes. Both pericyte coverage and numbers were normalized to the average of the control groups for each tamoxifen dose.

For vessel parameter analysis, images from the tomato lectin-stained channel were exported to ImageJ. MIP images were then generated and converted to 8-bit grayscale. A Gaussian blur (σ = 2) was applied, followed by triangle thresholding to delineate vascular structures. A binary mask was created, excluding background elements smaller than 20 square pixels to isolate the stained vessels. The resultant binary images, as illustrated in Supplementary Figure S1, were imported into AngioTool software for quantitative vascular analysis. The parameters assessed included Vessel Percentage Area, Junction Density, and Average Vessel Length, providing comprehensive metrics of the vascular network.

### Real-Time Quantitative Polymerase Chain Reaction (RT-qPCR)

Snap-frozen brain tissues were processed for RNA isolation using the RNA Miniprep Plus Kit (Zymo Research, Catalog #R1055) following the manufacturer’s protocol. From the isolated RNA, 500 ng was used as a template for cDNA synthesis through reverse transcription with random primers. Quantitative polymerase chain reaction (RT-qPCR) analyses were subsequently performed using the miScript SYBR® Green PCR Kit (Qiagen, Catalog #218073) on the LightCycler 480 System (Roche Diagnostics). GAPDH cDNA served as an internal control to normalize expression levels. The relative expression levels of the target genes were determined by calculating the difference in cycle threshold (ΔCT) values between GAPDH and the target mRNA, followed by quantification using the 2^(-ΔCT) method. Each assay was conducted in triplicate and repeated across three independent experiments to ensure reliability and reproducibility.

### Behavioral tests

Y maze spontaneous alternation test was used to assess the spatial recognition memory of the mice. Y-maze apparatus consisted of arms each measuring 35 cm in length, 20 cm in depth, and 6 cm in width, with a white finish. Each mouse’s behavior was observed and recorded for 5 minutes following its release into the maze. To maintain environmental consistency and eliminate olfactory cues, the maze was thoroughly cleaned with a 10% ethanol solution and air-dried between sessions. Video recordings from the experiment were manually analyzed to track the frequency and pattern of arm entries. A ‘spontaneous alternation’ was defined as the sequence in which a mouse entered all three arms consecutively without re-entering a previously visited arm. The Y-maze performance score, reflecting the animal’s spatial working memory, was calculated by dividing the number of spontaneous alternations by the total number of arm entry triads. Mice with fewer than 5 total arm entries were excluded from the analysis due to insufficient mobility.The locomotor activity of the mice was evaluated using Kondziela’s inverted screen (grip strength) test. Each mouse was initially placed at the center of a 30×30 cm screen made of 1 mm squared mesh. To begin the test, the screen was gently inverted, encouraging the mouse to grip the mesh to prevent falling. This procedure was repeated three times per mouse, and the duration of the grip was recorded either until the mouse fell or until a maximum time of 90 seconds was reached. The endurance time, defined as the length of time each mouse held onto the inverted screen before falling, was recorded for each trial. The average holding time was then calculated and compared between the experimental and control groups to assess differences in locomotor capabilities.

### Statistical Analysis

All statistical analysis was performed by GraphPad Prism 10 (version 10.2.0). Survival analysis was conducted using the Log-rank (Mantel-Cox) test. The difference between the body weight between groups were analyzed using two-way ANOVA followed by Sidak’s post hoc tests. Student’s t-test was used to compare pericyte coverage, count, vascular parameters, and motor and memory functions between groups. P values less than 0.05 were considered statistically significant.

## RESULTS

### Acute ablation of pericytes results in dose-dependent changes in body weight and survival

To create an inducible pericyte ablation model, we crossbred two transgenic mouse lines: *PDGFRβ-P2A-CreER*^*+/-*^ and *Rosa26-DTA176*^*+/+*^. The *PDGFRβ-P2A-CreER*^*T2*^ line expresses the *CreER*^*T2*^ complex under the *Pdgfrβ* promoter, which activates only upon tamoxifen administration. This activation facilitates the excision of *loxP*-flanked sequences in *Pdgfrβ* expressing cells. As a mating partner to this line, the *Rosa26-DTA176* line contains a *loxP*-flanked stop codon sequence upstream of the diphtheria toxin A (*DTA*) transgene, controlled by the constitutive *ROSA26* promoter. Tamoxifen-induced Cre recombinase excises the stop codon sequence, allowing DTA expression specifically in PDGFRβ^+^ cells. This strategy allowed us to selectively ablate PDGFRβ-expressing pericytes in the progeny, with CreER^+^ progeny serving as the pericyte ablation group upon tamoxifen administration, while CreER^-^ mice served as controls in characterization experiments (Figure 1a).

**Figure 1.**
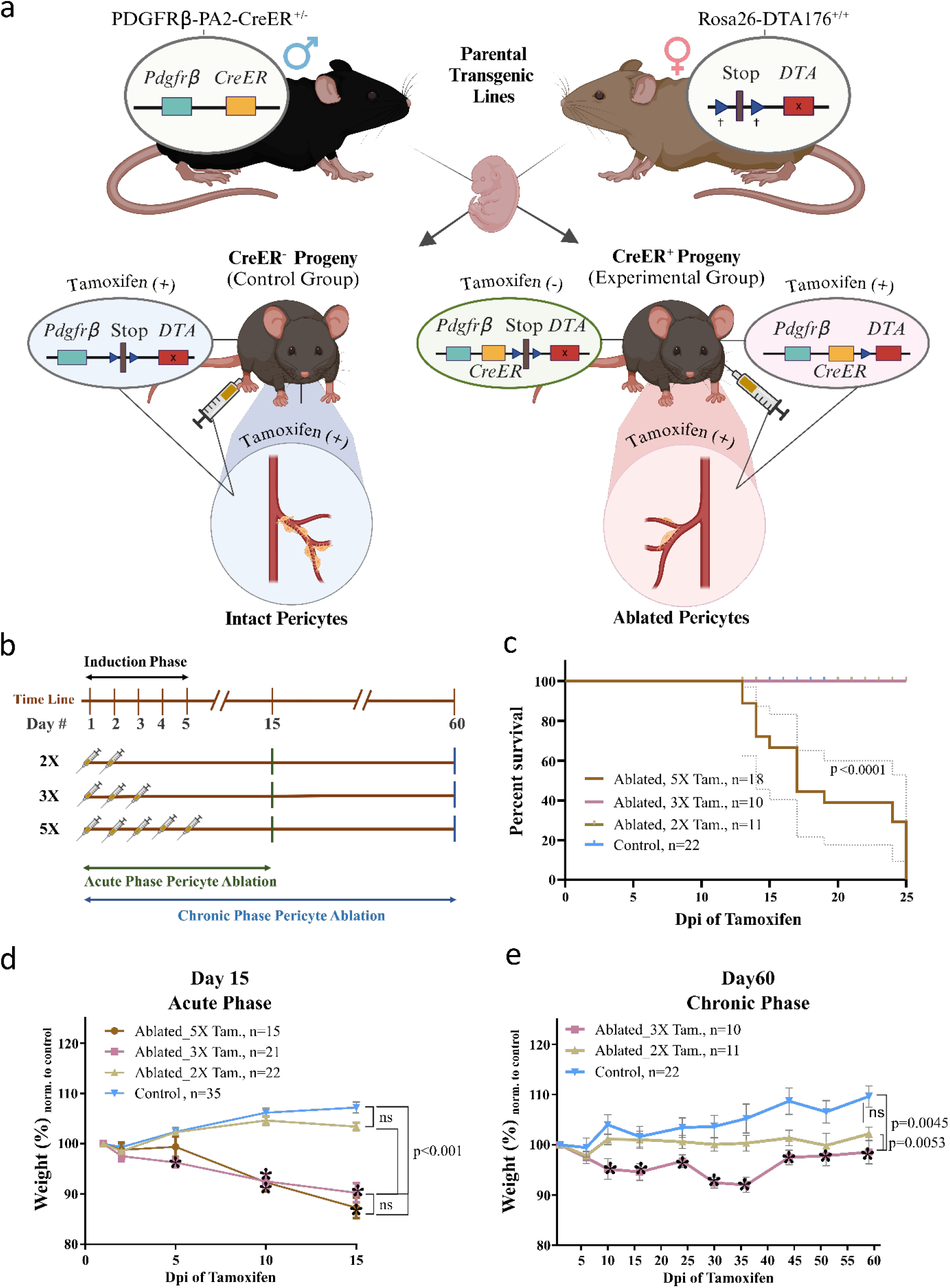
Generation of a Temporally Controlled Pericyte Ablation Model in Mice and the impact of pericyte ablation on body weight. **(a)** Diagram illustrating the generation of a tamoxifen-inducible and cell type-specific pericyte ablation model through the crossbreeding of *PDGFRβ-P2A-CreER*^*+/-*^ and *Rosa26-DTA176*^*+/+*^ transgenic mouse lines. CreER^+^ progeny was used as the pericyte ablation group, and CreER^-^ progeny served as a control group. Both groups received tamoxifen injections depending on their repetitive dosage group. Created by Biorender.com. **(b)** Schematic of the tamoxifen induction protocol depicting dosage schedules (2X, 3X, and 5X) and subsequent evaluation time points (Day 15 for acute phase and Day 60 for chronic phase). **(c)** Kaplan-Meier survival analysis of mice subjected to varying tamoxifen dosages, highlighting the survival impact of post-pericyte ablation over 25 days. Censorship is indicated by tick marks; p-values calculated using Log-rank (Mantel-Cox) test, noted as <0.0001. Error bars denote 95% confidence intervals. **(d-e)** Graphs depicting body weight trajectories of control versus pericyte-ablated mice during the acute (15 days) **(d)** and chronic (60 days) **(e)** phases post-tamoxifen induction. Data are presented as mean ± SEM for n=10-24 mice per group. Statistical analysis was conducted using two-way repeated measures ANOVA followed by Sidak’s post hoc test, with p<0.05 considered significant. No significant differences in body weight were detected between the CreER-control groups treated with 2X, 3X, and 5X tamoxifen; thus, these groups were combined for statistical comparison. In the acute phase **(d)**, significant weight differences were observed between in the 3X and 5X CreER^+^ groups compared to controls. In the chronic phase **(e)** there was a statistically significant difference between 3X CreER^+^ mice compared to controls. Asterisks denote the statistically significant results of post hoc multiple comparisons conducted to assess weight differences at individual time points compared to the CreER^-^ controls. **X=**Repetitive doses of tamoxifen injection, **Tam. =** Tamoxifen, **Dpi=** day post-injection, blue box=Pdgfrβ, **brown box=** translational stop sequence, **blue arrows (†)=**loxP sites, red box with **“x”=** untranslated DTA, **red box=** translated DTA, **yellow box=**Cre recombinase fused with estrogen receptor.

We first investigated the effect of varying tamoxifen doses to identify the optimal dosage by administering 100 mg/kg/day tamoxifen for 2, 3, or 5 consecutive days (2X, 3X, and 5X groups) by assessing both acute (Day 15) and chronic (Day 60) outcomes (Figure 1b). Notably, survival analysis demonstrated a significant impact of the highest tamoxifen dose (5X) on the survival of mice (p < 0.0001) (Figure 1c). Mice in the 5X group started to have a sick phenotype 12 days postinjection and none of them survived beyond three weeks (supplementary figure 2), whereas all animals in the 2X and 3X groups remained viable throughout the 60-day observation period. utopsies could be performed on a few mice immediately after their death and revealed large, swollen, dark-colored intestines (supplementary figure 3). As all animals in the 5X group died until Day 25, we could make examinations only in the acute phase for this group.

There were significant body weight differences between pericyte-ablated and CreER^-^ control mice in the 3X and 5X tamoxifen groups in the acute phase, but not in the 2X group (Figure 1d). In the 3X tamoxifen group, weight loss was evident on Day 4 and became markedly significant by Day 7 (p < 0.0001), culminating in a 17% difference by Day 15 (Figure 1d). Similarly, the 5X tamoxifen group showed statistically significant weight loss starting from Day 7, reaching a 20% reduction by Day 15 (p < 0.0001).

Long-term monitoring over 60 days revealed that the 3X tamoxifen group maintained lower body weight compared to controls, although a partial weight recovery was observed around 40 days post-tamoxifen injection (Figure 1e). In contrast, the 2X tamoxifen group showed no significant weight differences throughout the observation period. The data suggest a dose-dependent effect of tamoxifen on survival and body weight in CreER^+^ pericyte-ablated mice.

### Tamoxifen-induced ablation reveals regional variability and complex recovery dynamics of pericytes throughout the CNS

Immunofluorescence analysis was conducted to assess the degree of pericyte ablation in both cortical and spinal regions of mice after various repetitive doses of tamoxifen induction at acute and chronic phases (Figure 2). Quantitative analysis showed an effective reduction in both pericyte coverage and counts in both cortical and spinal cord tissues for each repetitive dosage (Figure 2 and Figure 3). In the acute phase, pericyte coverage in the cortex decreased by 83%, 81%, and 75% in the 2X, 3X, and 5X tamoxifen-treated mice, respectively, compared to controls (Figure 2a and 2c). In the chronic phase, cortical pericyte coverage was reduced by 67% and 71% compared to controls for the 2X and 3X doses, respectively. (Figure 2b and 2c). The analysis of pericyte count in the cortex showed a 58% (in 2X group), 44% (in 3X group), and 55% (in 5X group) decrease compared to controls during the acute phase, and 31% (in 2X group) and 48% (in 3X group) decrease compared to controls during the chronic phase (Figure 2d). The effectiveness of the ablation was also confirmed in the mRNA level with the target gene *PdgfrB* (Supplementary Figure 4).

**Figure 2:**
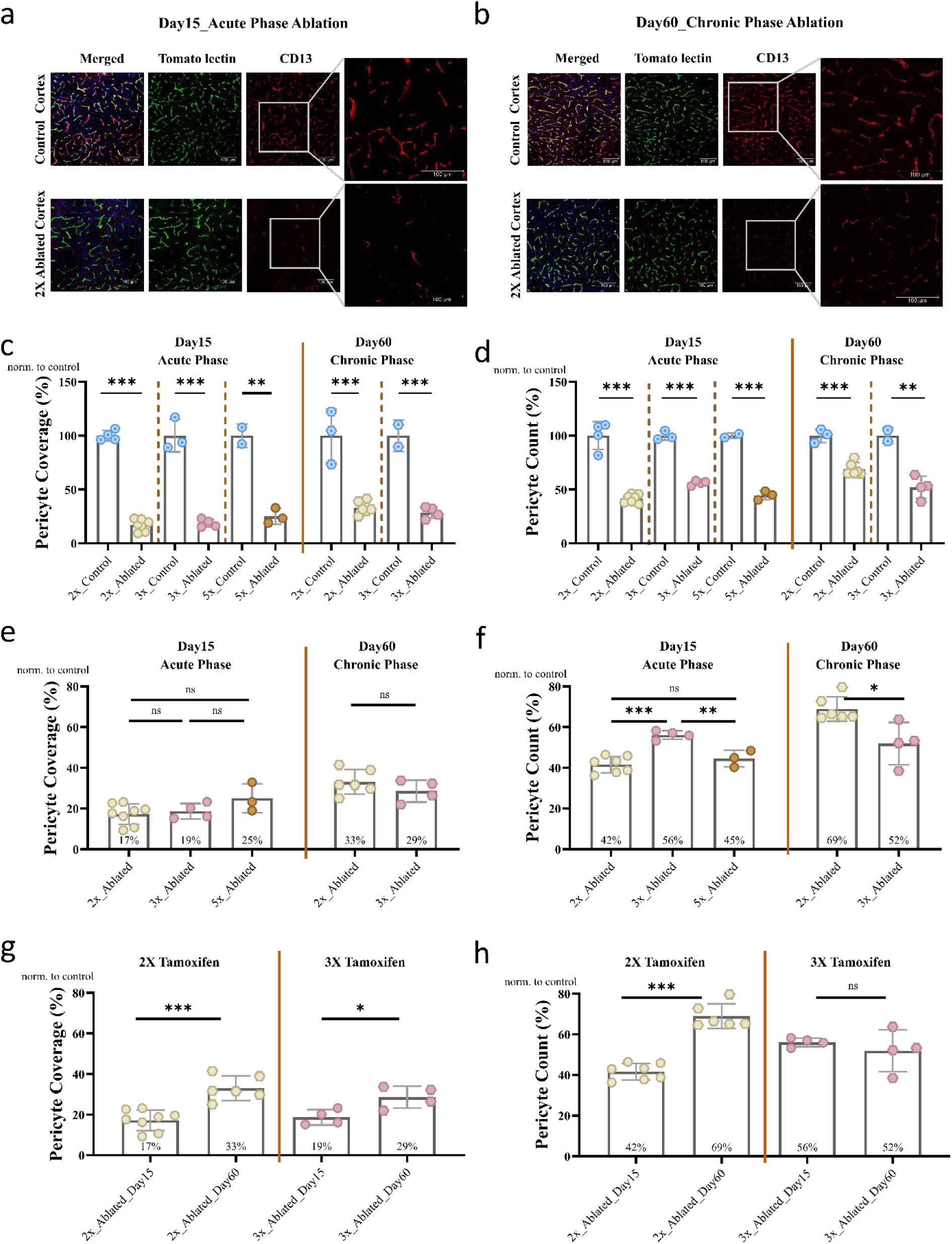
Immunofluorescent Characterization of Pericyte Ablation in Murine Cortex. **(a, b)** Immunofluorescence images showcasing pericytes (CD13, red), blood vessels (Tomatolectin, green), and nuclei (Hoechst, blue) within the cortex at acute (Day 15) and chronic (Day 60) phases post-tamoxifen induction, evidencing significant pericyte loss. **(c-h)** Quantitative analysis of pericyte coverage **(e, g)** and count **(f, h)**, demonstrating dose-dependent **(e, f)** and time-course effects **(g, h)** of tamoxifen-induced ablation. Statistical significance determined by Student’s t-test, with p-value annotations: ns (0.1234), * (0.0332), ** (0.0021), *** (0.0002). Data points reflect averages of 6-8 cortical images per animal. **X =** repetitive doses of tamoxifen injection.

**Figure 3:**
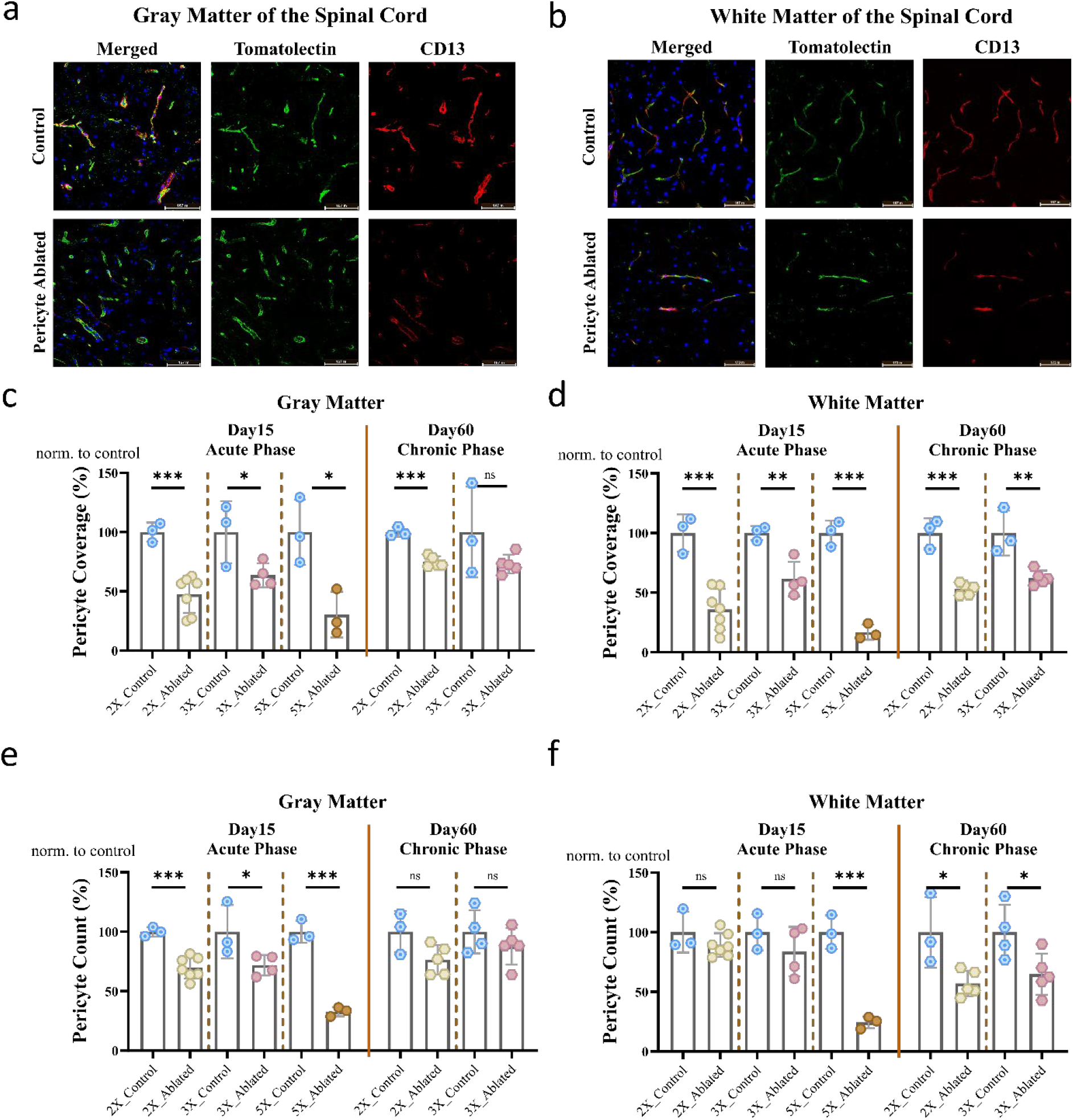
Region-Specific Effects of Pericyte Ablation in Spinal Cord Compared to Controls. **(a, b)** Immunofluorescent staining of spinal cord sections (gray matter and white matter, respectively) post 2X tamoxifen induction, revealing stark pericyte coverage reduction. **(c-f)** Histograms quantifying pericyte coverage and count in gray and white matter, after acute and chronic phases of pericyte ablation. Statistical analysis via Student’s t-test, p-values classified as ns (0.1234), * (0.0332), ** (0.0021), *** (0.0002). Data points represent the averages based on 8-10 images from various spinal cord segments per animal. **X =** repetitive doses of tamoxifen injection.

When comparing the effects of different tamoxifen doses on pericyte coverage, no significant differences were observed across the dosages during either the acute or chronic phase (Figure 2e). Notably, despite the decreased survival following the 5X dose, pericyte numbers in the cortex were comparable to those seen with the 2X dose. Interestingly, the 3X dose caused a less pronounced reduction in pericyte count in the acute phase compared to both the 5X and 2X doses. However, in the chronic phase, the normalized pericyte count was significantly lower in the 3X group compared to the 2X dose, indicating a more prolonged effect of the 3X tamoxifen dose (Figure 2f).

The comparison of pericyte ablation in the cortex between the acute and chronic phases revealed a recovery in both pericyte coverage and numbers during the chronic phase. This recovery was most notable in the 2X dose group, where pericyte numbers increased from 42% to 69% of the control group, and pericyte coverage improved from 17% to 33% (Figure 2g and 2h). The 3X group also showed some recovery, with coverage increasing from 19% in the acute phase to 29% in the chronic phase (Figure 2g). However, pericyte numbers remained stable in the 3X dose group, suggesting a diminished capacity for replenishment as the tamoxifen dose increases (Figure 2h). These findings indicate that CNS pericytes have the ability to regenerate over time following acute ablation.

Analysis of pericytes in the spinal cord confirmed the effectiveness of our pericyte ablation model in this CNS region as well (Figures 3a and 3b). In the acute phase, pericyte coverage in the gray matter was reduced to 47%, 64%, and 30% of control levels following 2X, 3X, and 5X tamoxifen doses, respectively (Figure 3c). Similarly, in the white matter, pericyte coverage decreased to 36%, 62%, and 17% of control levels for the 2X, 3X, and 5X doses, respectively (Figure 3d). Sixty days after tamoxifen administration, pericyte coverage remained reduced in both the gray and white matter of the 2X group and the white matter of the 3X group compared to controls (Figures 3c and 3d).

Pericyte numbers were also decreased in the spinal cord after tamoxifen administration (Figures 3e and f). On day 15, this decrease was evident only in the gray matter, but not in white matter, of the 2X and 3X tamoxifen groups, whereas pericyte count was lower in both the gray and white matter of the 5X tamoxifen group compared to controls (Figures 3e and 3f). On day 60, there was a trend for decreased pericyte counts in the gray matter of the 2X tamoxifen group (Figure 3e). In the white matter, both 2X and 3X tamoxifen groups had lower pericyte counts compared to controls, indicating that tamoxifen-induced pericyte loss occurs later in the white matter of the spinal cord compared to the gray matter and the cortex. (Figure 3f).

When we compared the effect of different doses, the 5X dose resulted in a greater reduction in pericyte coverage and numbers compared to 2X and 3X doses in the spinal cord gray matter (Figures 4a and 4c). In the white matter, 2X and 5X dosages induced a significantly greater reduction in pericyte coverage compared to the 3X tamoxifen dose (Figure 4b), and the 5X dose resulted in a greater reduction in pericyte numbers compared to 2X and 3X doses (Figure 4d). Next, we evaluated the capacity for pericyte coverage and the number of recoveries in the spinal cord following acute ablation. Sixty days after tamoxifen induction, we observed an increase in pericyte coverage in the gray matter, with a similar upward trend in the white matter in the 2X tamoxifen group compared to Day 15 (Figure 4e-f). In contrast, pericyte coverage in the 3X tamoxifen group remained unchanged in both the gray and white matter when compared to Day 15. No change was detected in the gray matter for pericyte counts, while a decrease was noted in the white matter of the 2X tamoxifen group during the chronic phase compared to the acute phase (Figure 4g-h). No statistically significant changes in pericyte counts were observed in the 3X group during the chronic phase.

These results indicate that the extent of pericyte loss is greater in the cortex than in the spinal cord (Supplementary Figure 5), with more prolonged pericyte death observed in the white matter of the spinal cord compared to the gray matter. Additionally, cortical pericytes exhibit a higher recovery rate than those in the spinal cord.

**Figure 4:**
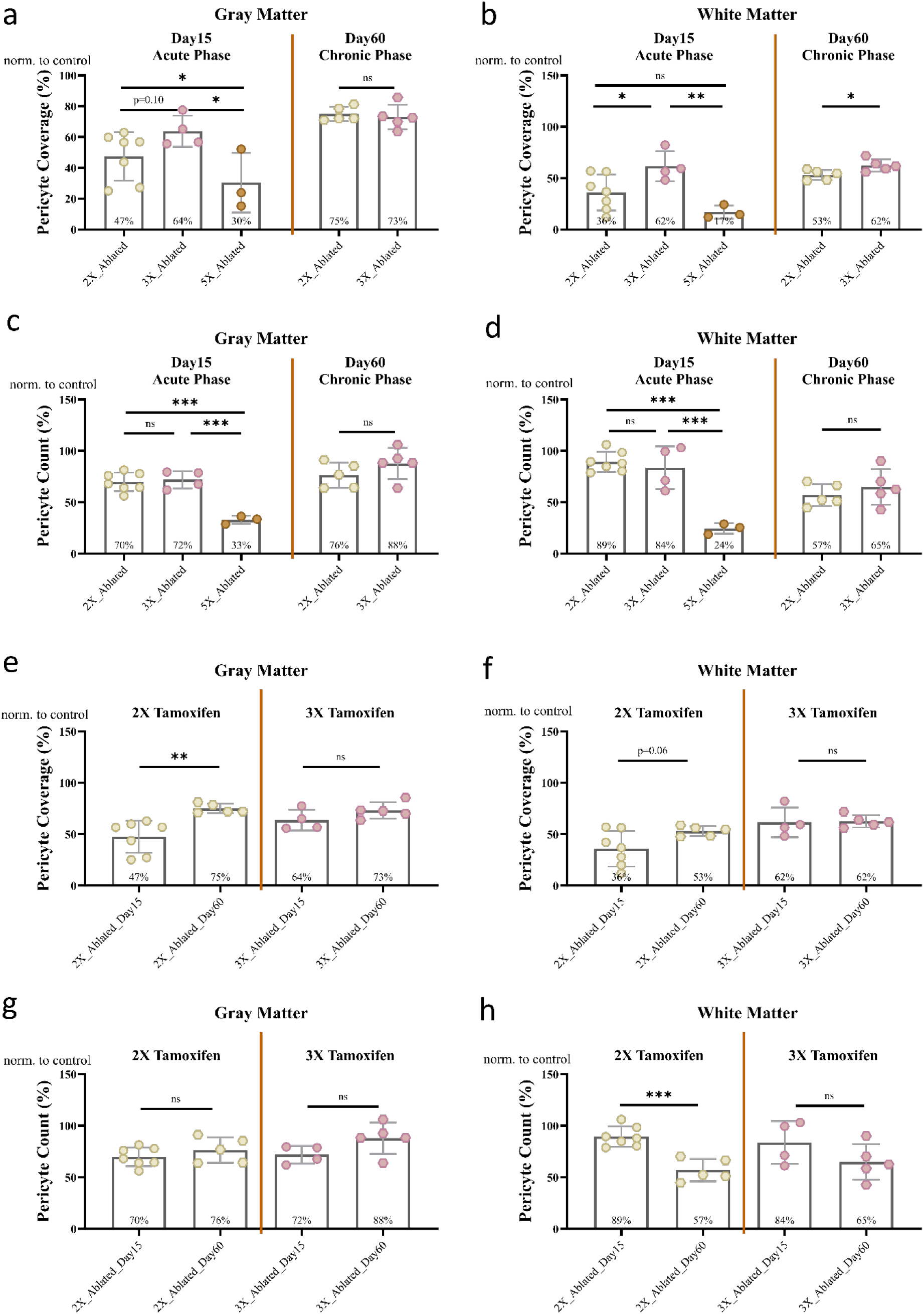
Pericyte Analysis in the Spinal Cord Across Tamoxifen Doses. **(a, b, c, d)** Comparison of pericyte coverage **(a, b)** and pericyte numbers **(c, d)** in gray matter **(a, c)** and white matter **(b, d)** at acute (Day 15) and chronic (Day 60) phases post-tamoxifen across all tamoxifen doses. **(e, f, g, h)** Comparisons of pericyte coverage **(e, f)** and pericyte number **(g, h)** in acute and chronic phases for 2X and 3X tamoxifen doses in gray matter **(e, g)** and white matter **(f, h)**. Statistical significance was assessed as ns (0.1234), * (0.0332), ** (0.0021), *** (0.0002). Data is taken from multiple spinal cord sections per animal. X = repetitive tamoxifen doses. Data points represent the averages based on 8-10 images from various spinal cord segments per animal. X = repetitive doses of tamoxifen injection.

### Vascular parameters are not changed after acute pericyte ablation

Vessel Percentage Area (VPA), Junction Density (JD), and Average Vessel Length (AVL) were assessed in the cortex and spinal cord in the acute and chronic phases following 2X tamoxifen administration. There was no statistically significant change in VPA, JD, and AVL in the cortex (Figure 5a-c) and the spinal cord (Figure 5d-i). These findings indicate that inducible pericyte ablation does not significantly alter vascular morphology for up to 60 days in CNS.

**Figure 5:**
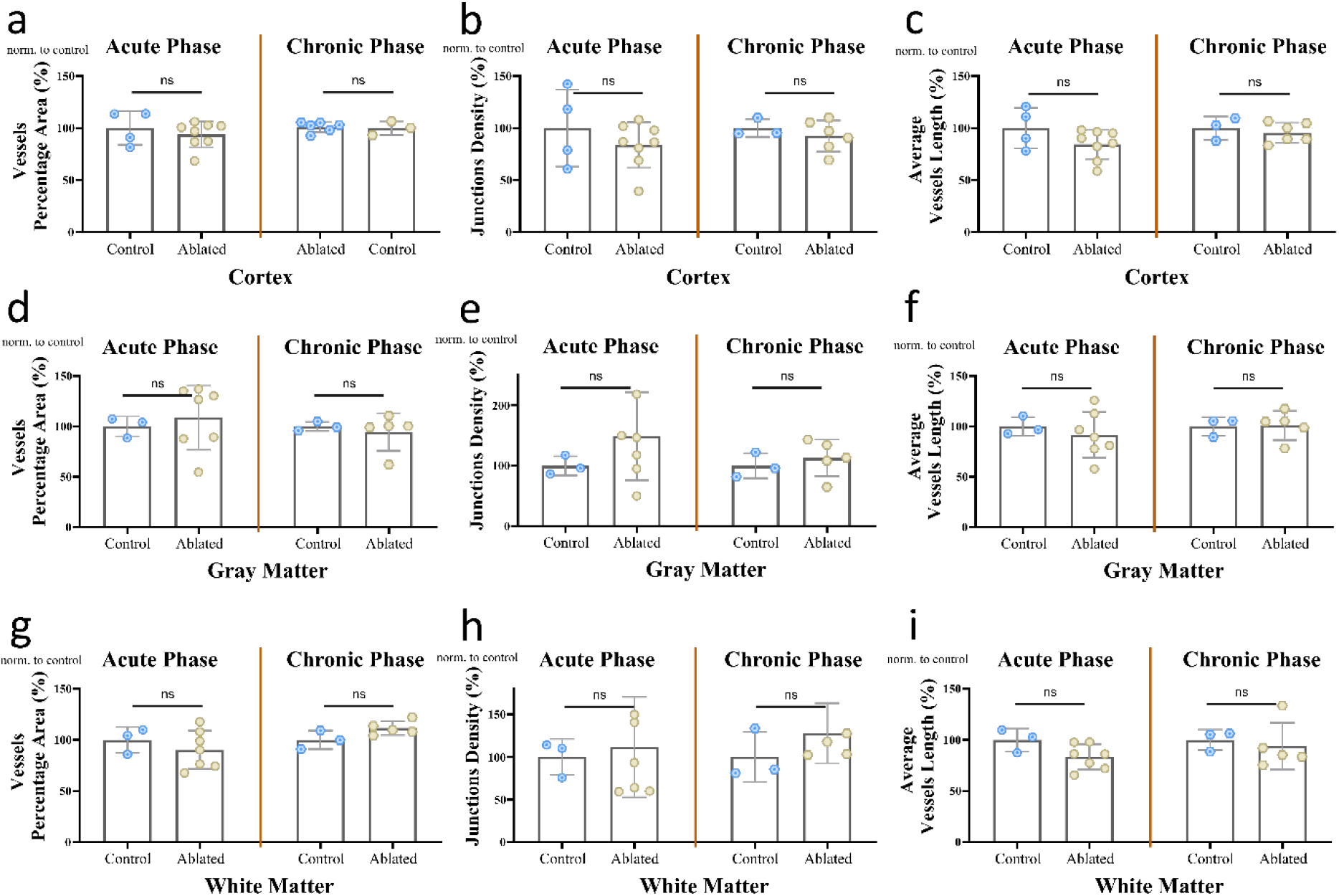
Vascular Parameter Quantification Post-Pericyte Ablation in Cortex and Spinal Cord. A comprehensive evaluation of vascular parameters: **(a, d, g)** Vessel Percentage Area [VPA], **(b, e, h)** Junction Density [JD], and **(c, f, i)** Average Vessel Length [AVL] in the **(a-c)** cortex and **(h-i)** spinal cord during acute and chronic phases post-ablation with 2 repetitive doses of Tamoxifen induction. Statistical analysis was performed using Student’s t-test; significance denoted as ns (not significant, p=0.1234), * (p=0.0332), ** (p=0.0021), *** (p=0.0002). Data aggregated from 6-8 (cortex) and 10-12 (spinal cord) randomized images per animal. Two-way ANOVA with Bonferroni’s correction applied for statistical analysis, setting significance at p < 0.05. Asterisks indicate significant differences.

### Pericyte Ablation in Adult Mice Does Not Impair Spatial Memory or Muscle Strength

Using the Y-maze spontaneous alternation task, we assessed motor function using Kondziela’s inverted screen test and spatial memory. In Kondziela’s inverted screen test, the time it took for the mice to fall was similar to controls across all groups, both in the acute and chronic phases (Figure 6a). No significant differences were observed when comparing different tamoxifen doses or time points (Figure 2c and Figure 2e). Similarly, in the Y-maze spontaneous alternation test, pericyte ablation did not significantly affect performance in any group during either the acute or chronic phases (Figure 6b). Comparisons across different doses and time points also revealed no significant differences (Figure 6d and Figure 6f).

**Figure 6:**
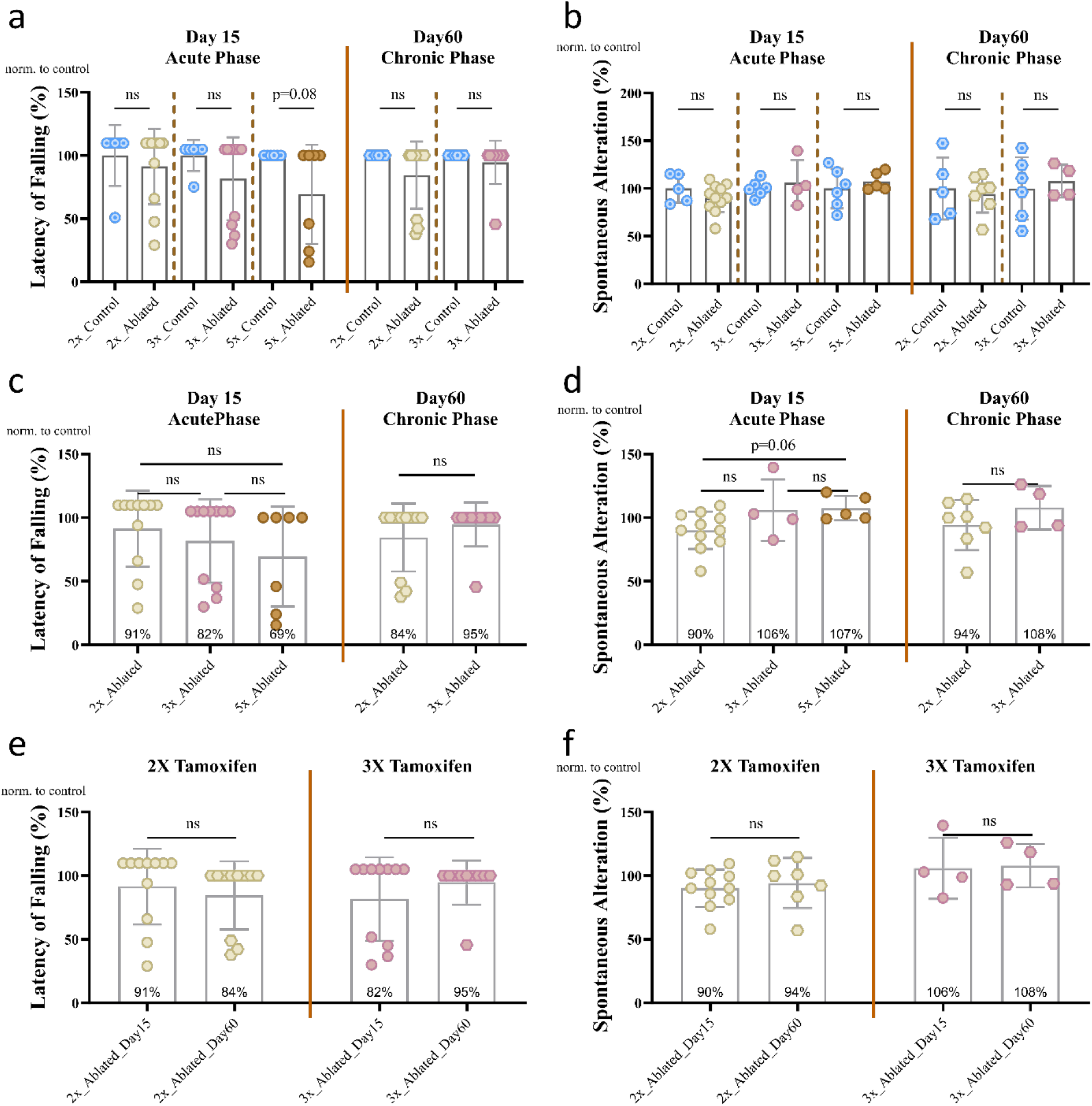
Motor and Cognitive Functional Assessment Following Pericyte Ablation. **(a, b)** Motor coordination and cognitive function were assessed using the inverted screen test and Y maze spontaneous alternation task, respectively, in mice following tamoxifen-induced pericyte ablation. Results across acute and chronic phases are compared for control versus pericyte-ablated groups at tamoxifen dosages of 2X, 3X, and 5X. **(c, d)** Analysis in the chronic phase, incorporating both motor coordination and cognitive assessments, further evaluates the effects of varying tamoxifen doses, highlighting dose-dependent impacts on function. **(e, f)** Longitudinal comparison of motor and cognitive performance, specifically contrasting the effects of 2 and 3 tamoxifen doses across acute and chronic phase. Statistical relevance assessed via Student’s t-test, categorizing significance as not significant (ns, p=0.1234), * (p=0.0332), ** (p=0.0021), *** (p=0.0002), with data presented as mean ± SEM.

## DISCUSSION

In this study, we present a novel tamoxifen-inducible acute pericyte ablation model based on the induction of an attenuated diphtheria toxin variant (DTA176) in *Pdgfrβ*-expressing cells. Using this model, we have demonstrated that a precise tamoxifen dosage adjustment can achieve substantial pericyte depletion within the CNS without adversely affecting animal health. Additionally, we observed that pericytes regenerate *in vivo*, and pericyte coverage is restored in the subsequent weeks following extensive pericyte loss, reinforcing the concept of pericyte plasticity. Moreover, our findings indicate that the extent of pericyte ablation differs between the brain and spinal cord, suggesting regional heterogeneity of pericytes within the CNS.

The model proposed here is based on two transgenic strains: *PDGFRβ-P2A-CreER*^*T2*^ and *Rosa26-DTA176*. The *PDGFRβ-P2A-CreER*^*T2*^ strain, developed by Cuervo et al.^35^, has been previously utilized to study pericytes in the retina, brain, and lung^37-39^. By crossing with *Rosa-tdTomato* mice, Cuervo et al. demonstrated tdTomato reporter expression in 84% of NG2-expressing pericytes within the retinal vasculature. Our study observed a 75-83% reduction in cortical pericytes, indicating that DTA176 induction occurred in approximately 80% of pericytes, consistent with Cuervo et al.’s findings.

Previously, Eilken et al. developed the *Pdgfr-β-CreER*^*T2*^ mice^40^, which they crossed with *ROSA-DTA* mice^41^ to create a tamoxifen-inducible acute pericyte ablation model (*DTA*^iPC^). This model resulted in severe weight loss postnatally, with animals failing to survive beyond one week. To mitigate this, Eilken et al. utilized the *ROSA26-iDTR*^*42*^, which expresses the diphtheria toxin receptor post-tamoxifen administration and requires subsequent diphtheria toxin injection for pericyte ablation (*DTR*^iPC^). In our model, we observed weight loss with 3X and 5X tamoxifen doses but not with the 2X dose. Furthermore, we noted reduced survival rates two weeks post-5X tamoxifen administration, an effect absent with lower doses. These observations suggest that acute ablation of *Pdgfrβ*-expressing cells may precipitate severe systemic effects. Therefore, this model necessitates stringent tamoxifen dosage control to avoid off-target effects on other *Pdgfrβ*-expressing cells. As the 2X tamoxifen dose did not induce weight loss and the animals exhibited no complications or behavioral changes for at least two months, this regimen can be ideal for future pericyte ablation studies.

Cornuault et al. recently employed *DTA*^iPC^ mice to investigate pericyte ablation effects on cardiac function in 8-week-old mice^43^.

Tamoxifen was administered (50 mg/kg/day) for five consecutive days, and the dose was repeated biweekly to sustain stable pericyte ablation (60% capillary pericyte depletion after two injection series), indicating robust renewal potential of cardiac pericytes. In our study, pericyte renewal was also observed after the lowest tamoxifen dose within the brain (increasing from 42% to 69% of controls in the cortex), highlighting the renewal potential of CNS pericytes. Similar to Eilken et al., Cornuault et al. reported significant weight loss in pericyte-ablated mice and their systemic analysis revealed near-total depletion of pericytes in the aorta media, intestinal lacteals, and skeletal muscles, with a significant inverse correlation between the degree of cardiac pericyte depletion and weight loss. Consistently, we also observed significant weight loss, severe intestinal pathology and decreased survival with the 5X tamoxifen dosage. These results demonstrate that tamoxifen dose must be carefully adjusted and lower doses should be administered to avoid serious systemic affects in *Pdgfrβ-* dependent acute pericyte ablation models.

To circumvent the systemic side effects associated with *Pdgfrβ*-expressing cell ablation, Nikolakopoulou et al. devised a pericyte-specific ablation model utilizing a double-promoter approach with *Pdgfrβ* and *Cspg4*^31^. In this model, pericyte-specific Cre mice are crossed with *iDTR* mice, and pericyte ablation is induced by seven days of tamoxifen administration (40 mg/kg/day) followed by ten days of diphtheria toxin administration (0.1 μg/day) commencing two weeks post-tamoxifen. This model achieved 60% pericyte ablation in the cortex 15 days after the final injection. However, this model’s limitations include the requirement for repeated diphtheria toxin injections that can be related to potential off-target effects, and limited commercial availability of double-promoter mice.

Another tamoxifen-inducible pericyte ablation strategy, as employed by Vazquez-Liebanas et al., involves the conditional knockout of *Pdgfβ* from *Cdh5*-expressing endothelial cells^34^. In this model, *Pdgfb* deletion in 2-month-old mice induces gradual pericyte loss, resulting in a 50% reduction in pericytes by 12-18 months of age. Unlike *Pdgfrβ*-or double-promoter-based models, pericyte loss in this model is not acute and exhibits less controllability.

Previously, using the double-promoter mice, Kisler et al. demonstrated that cortical vascular density remains unchanged three days following pericyte ablation in adult mice^20^. Consistent with this finding, we found no significant alterations in vascular percentage area, average vessel length, or junction density in the cortex and spinal cord 15 days post-pericyte ablation. Furthermore, we observed that brain vascular morphology is preserved up to 60 days post-induction, despite sustained pericyte loss. Similarly, using the inducible *Pdgfb* knockout model, Vazquez-Liebanas et al. found that adult-induced pericyte loss does not result in vessel dilation, impaired arteriovenous zonation, or microvascular calcifications, contrary to the effects seen with constitutive *Pdgfb* loss^25,44^. Collectively, these findings suggest that while pericytes are essential for prenatal vascular development, they are not critical for the maintenance of the mature vascular network in the nervous system during adulthood.

We observed a lower reduction in pericyte numbers and coverage following tamoxifen administration in the spinal cord gray and white matter compared to the cortex. Pericyte coverage and number in the spinal cord were previously quantified by Winkler et al.^21^. The authors reported that PDGFRβ+ pericyte coverage was approximately 80% and pericyte density was around 2000/mm^2^ in the cortex. In the spinal cord anterior horn, both pericyte coverage (∼55%) and density (∼1100/mm^2^) were lower compared to the brain, and in the spinal cord white matter (∼70% and ∼1800/mm^2^, respectively). In the same study, *Pdgfrβ*^*F7/F7*^ mice^28^ were used to assess the extent of spinal cord pericyte ablation, revealing a ∼40% reduction in the anterior horn, which was less pronounced compared to the 60% loss observed in the cortex^22^. This suggests that the reduced degree of pericyte ablation observed in our study in the spinal cord may be attributed to the lower baseline presence of pericytes in this region. On the other hand, Göritz et al. identified two types of pericytes in the spinal cord and brain, one of which contributes to fibrosis post-injury^17,18^. Similarly, Birbrair et al. identified two types of pericytes (Nestin-negative and -positive) across various organs, including the brain and spinal cord^45^. These studies indicate molecular heterogeneity among pericytes in different CNS regions; therefore, it is possible that *Pdgfrβ* expression may be lower in spinal cord pericytes. Future studies are required for a more precise explanation for this aspect.

The temporal characteristics of pericyte coverage and count following acute pericyte ablation in adult mice have not been investigated before. In our study, we observed that a lower tamoxifen dosage (2X) facilitated partial recovery of both pericyte count and coverage in the cortex during the chronic phase. Conversely, in mice administered a 3X tamoxifen dose, an increase in pericyte coverage, but not pericyte count, was noted. In the spinal cord gray matter, an increase in pericyte coverage, but not number, was observed only after the 2X dose. Previously, utilizing *PDGFRβ-Cre/YFP* mice and chronic cranial window imaging, Berthiaume et al. demonstrated that pericyte processes are dynamic under basal conditions and can extend to contact uncovered endothelial regions following the selective ablation of adjacent pericytes^46^. In a subsequent study, the same group reported that pericyte plasticity is compromised in aged mice^47^. Our findings indicate that CNS pericytes respond to ablation not only by extending processes to cover endothelial gaps but also by replenishing pericytes following more extensive ablation in vivo in adult mice, developing the concept of plasticity of CNS pericytes further. The origin of cells responsible for replenishing pericytes remains unknown and warrants further investigation.

There are some limitations to consider. In our model, we induce the ablation of *Pdgfrβ* expressing cells, which is not limited to pericytes and includes cells such as vascular smooth muscles cells. Additionally, pericyte ablation is not limited to CNS and the degree of pericyte ablation can be different in other organs, which should be studied in future studies. While previous research has demonstrated BBB leakage following induced pericyte ablation in adult mice, we did not investigate BBB integrity in this study^31,44^. Moreover, we did not focus on the exact mechanisms of pericyte recovery after the ablation, and further research is needed to elucidate these mechanisms.

In conclusion, our tamoxifen-inducible, cell-type-specific pericyte ablation model offers significant improvements in temporal control, specificity, and simplicity over existing models. It enables detailed studies of pericyte roles in adult CNS physiology without the developmental confounds of traditional models. However, dose-dependent toxicity and regional variability highlight the need for careful experimental design and interpretation. This model provides a robust platform for advancing our understanding of pericyte-mediated neurovascular regulation and developing targeted therapeutic strategies for neurovascular disorders.

## Author Contributions

DA contributed to data acquisition, analysis, interpretation, and preparation of the first draft of the manuscript. EY, AÖ, IAA, and ŞÜZ contributed to data analysis. EÖ contributed to data acquisition and analysis. MY, ABY, ABK, ÇK, and ZLC contributed to data acquisition. CİK and ET contributed to data interpretation and manuscript revision. YGÖ and AV conceived the study, and contributed to data analysis, interpretation, drafting, and revising of the manuscript.

## Acknowledgment

This study was conducted using the service and infrastructure of Koç University Research Center for Translational Medicine (KUTTAM).

