## Supplementary Data for "A Novel Mouse Model Demonstrates In Vivo Replenishment of Central Nervous System Pericytes After Successful Acute Ablation"

### 1 Supplementary Tables

**Table 1.** PCR Primers used for Genotyping of the progenies

| | |
|---|---|
| PDGFRβ-CreER and PDGFRβ WT Forward Primer | CCA CCT TGA ATG AAG TCA ACA C |
| PDGFRβ WT Reverse Primer | AGC TTG TGG CAG TGT AGC TG |
| PDGFRβ-CreER Mutant Reverse Primer | ACA TGT CCA TCA GGT TCT TGC |

**Table 2.** Touch-down PCR settings

| Step | Temperature ($^o$C) | Time (minutes) | Repeats and Delta Temperature settings |
|---|---|---|---|
| 1 | 94 | 2:00 | |
| 2 | 94 | 0:20 | |
| 3 | 65 | 0:15 | -0.5 C per cycle decrease |
| 4 | 68 | 0:10 | |
| 5 | | -- | repeat steps 2-4 for 10 cycles (Touchdown) |
| 6 | 94 | 0:15 | |
| 7 | 60 | 0:15 | |
| 8 | 72 | 0:10 | |
| 9 | | -- | repeat steps 6-8 for 28 cycles |
| 10 | 72 | 2:00 | |
| 11 | 10 | infinite | hold |

#### 2 Supplementary Figures

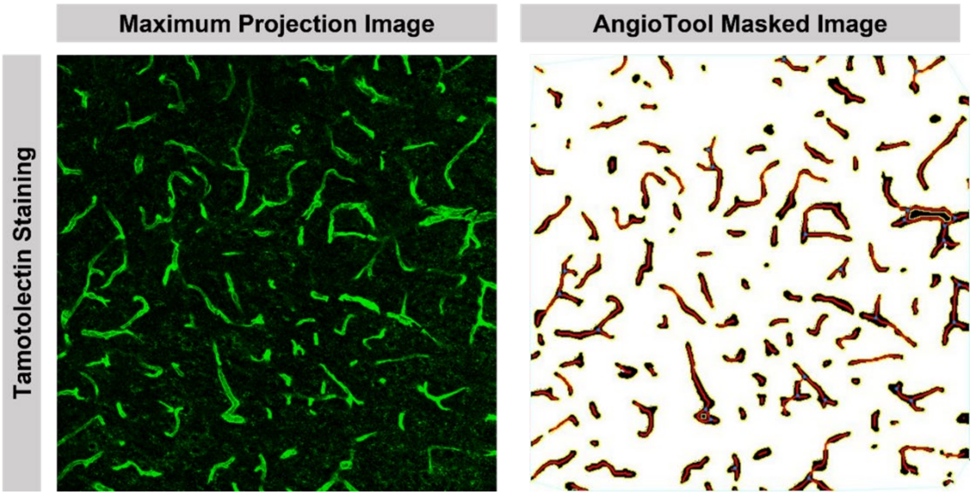


**Supplementary Figure 1**. The masked image generated by AngioTool Software for vessel analysis

AngioTool software was used to mask the Tomatolectin-stained vessels. Then, the mask images were analyzed in terms of vessel parameters.

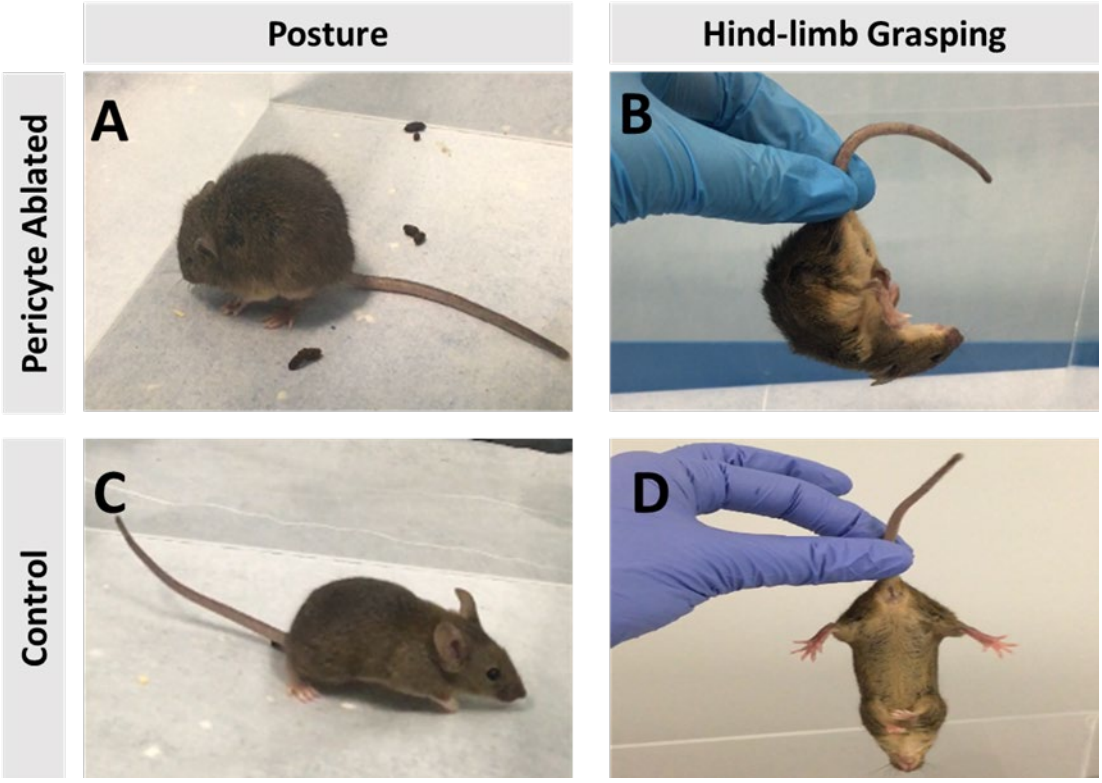


**Supplementary Figure 1**. The phenotypical changes after the pericyte ablation by 5X of tamoxifen administration

Postural change with the appearance of a humpback (a), and foot grasping reflex (b) was observed in the 5X tamoxifen given mice, whereas CreER$^{-}$ control mice had normal findings (c,d)

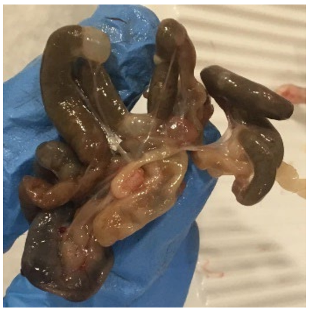

**Supplementary Figure 2.** Observations after the autopsy of the 5X tamoxifen group

Autopsy findings of an animal that died on day 20 postinjection revealed large, swollen, dark-colored intestines suggesting gastrointestinal infarction.

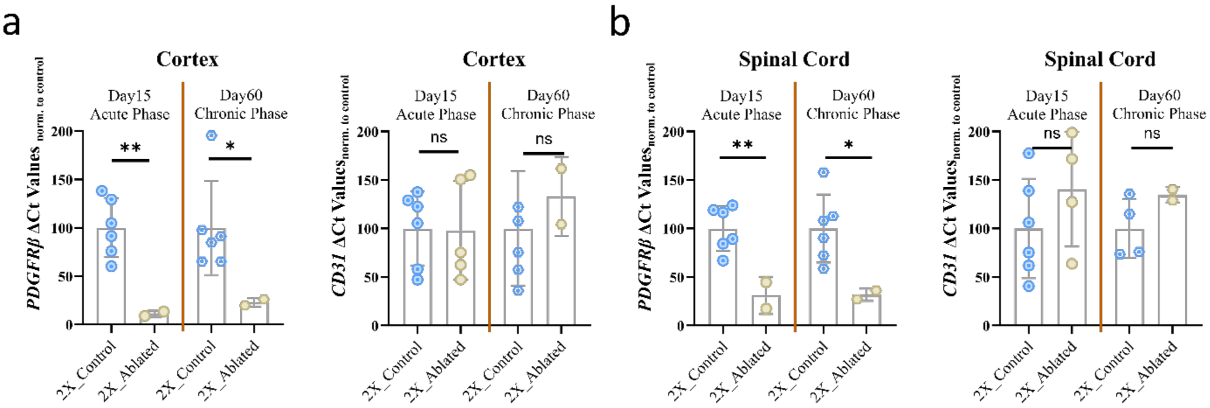


**Supplementary Figure 4.** Real-Time Quantitative PCR Results for Pdgfrβ and CD31 as Pericyte and Endothelial Markers

**(a, b)** Expression levels of Pdgfrβ and CD31 in the cortex at acute (Day 15) and chronic (Day 60) phases post-2X tamoxifen induction are shown as percentages relative to the control group. Pdgfrβ serves as a pericyte marker and. CD31 serves as an endothelial marker.

**(c, d)** Expression levels of Pdgfrβ and CD31 in the spinal cord at acute and chronic phases post-2X tamoxifen induction are shown as percentages relative to the control group.

Statistical analysis via Student's t-test, p-values classified as ns (0.1234), * (0.0332), ** (0.0021), *** (0.0002). Data points represent the averages based on 8-10 images from various spinal cord segments per animal. X = repetitive doses of tamoxifen injection.

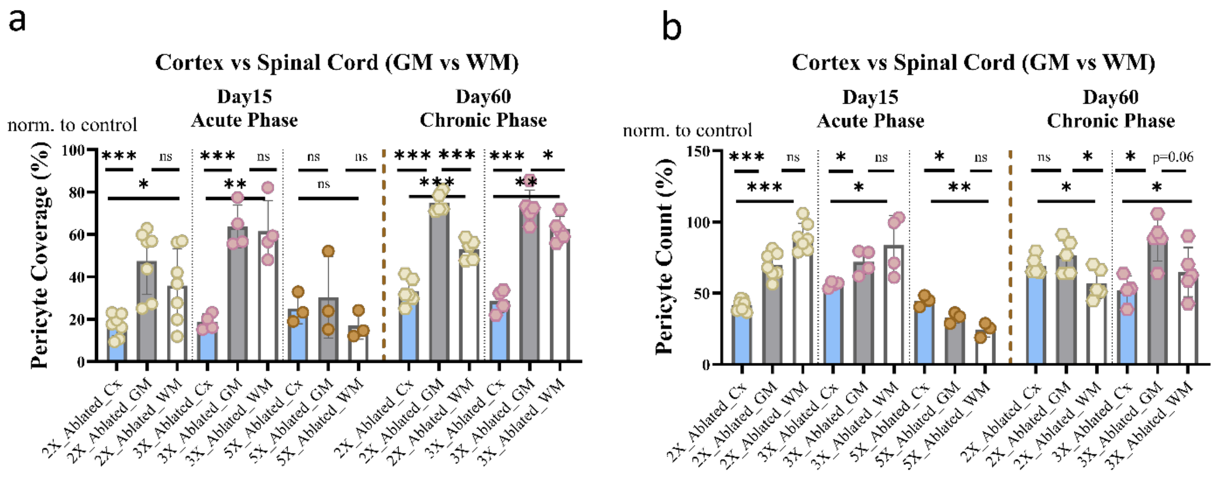


**Supplementary Figure 5.** Comparison of pericyte ablation in cortex, gray matter, and white matter of the spinal cord across tamoxifen dosages

**(a,b)** Comparison of the pericyte coverage **(a)** and pericyte numbers **(b)** in cortex, gray matter, and white matter in acute and chronic phases of pericyte ablation with different tamoxifen dosages. numb
